# Early-life adversity is associated with differential gene expression in response to acute psychological stress: preliminary findings

**DOI:** 10.1101/727180

**Authors:** Idan Shalev, Waylon J. Hastings, Laura Etzel, Salomon Israel, Michael A. Russell, Kelsie A. Hendrick, Megan Zinobile, Sue Rutherford Siegel

## Abstract

**Objective:** Exposure to early-life adversity (ELA) can result in long-term changes to physiological systems, which predispose individuals to negative health outcomes. This biological embedding of stress-responsive systems may operate via dysregulation of physiological resources in response to common stressors. The present study used a novel experimental design to test how young adults’ exposure to ELA influence neuroendocrine and inflammatory responses to acute stress.

**Materials and methods:** Participants were 12 males (mean age= 21.25), half of whom endorsed at least three significant adverse events up to age 18 years (‘ELA group’), and half who confirmed zero (‘controls’). Using a randomized within-subjects, between-groups experimental design, we induced acute psychosocial stress (Trier Social Stress Test, TSST), and included a no-stress control condition one week apart. During these sessions, we obtained repeated measurements of physiological reactivity, gene expression of *NR3C1, FKBP5* and *NFKB1*, and plasma levels of pro-inflammatory cytokines (IL-1β, IL-6, IL-8 and TNFα) over a 4-hour window post-test.

**Results:** The ELA group evinced significantly higher cortisol response and lower *NR3C1* gene expression in response to the TSST compared with controls, while no differences were observed in the no-stress condition. Cortisol and group status interacted such that increase in cortisol predicted increase in both *NR3C1* and *NFKB1* expression among controls, but decrease in the ELA group. For pro-inflammatory cytokines, only IL-6 increased significantly in response to the TSST, with no differences between the two groups.

**Conclusion:** Overall, we provide preliminary findings for the biological embedding of stress via a dynamic and dysregulated pattern evidenced in response to acute psychosocial stress. ELA may program physiological systems in a maladaptive manner more likely to manifest during times of duress, predisposing individuals to the negative health consequences of everyday stressors. Future studies with larger sample size including both males and females are needed to replicate these findings.

## Introduction

An ever-growing body of research suggests that early-life adversity (ELA) can program biological systems, which predispose individuals to later-life physical and mental-health problems [1, 2], Empirical evidence exist for associations between ELA and elevated risk of depression, cardiovascular disease, diabetes, autoimmune diseases and cancer, to name a few (reviewed in [3]). Despite the salient role of ELA on disease risk, the biological mechanisms that play a downstream role in increased disease susceptibility are not well understood.

Mechanistic research on the biological embedding of ELA has emphasized maladaptive programming of the hypothalamic-pituitary-adrenal (HPA) axis with the associate release of cortisol through processes of allostasis [3, 4]. Specifically, studies have documented a shift in HPA axis function with hyper-or hypo-secretion of cortisol in depression and post-traumatic stress disorder (PTSD), respectively [5, 6]. Similar findings have been reported in individuals exposed to ELA without such diagnoses [7, 8]. This programming, in turn, can result in mitochondrial dysfunction, failure to down-regulate the inflammatory response and overall metabolic stress, thereby increasing circulatory levels of lipids, glucose, oxidants, and pro-inflammatory cytokines [9, 10]. Further mechanistic research on the biological embedding of ELA suggests physiological dysregulation may be mediated at the genetic level via epigenetic modifications that can persist over long periods of time [11], including evidence linking ELA and cortisol responses via methylation levels in the glucocorticoid receptor *(NR3C1)* gene [12, 13]. Other research suggest the involvement of telomere biology in mediating the longer-term link between ELA and disease risk [14]. What is less clear, however, is how target immune cells respond to stress *in vivo* as a consequence of ELA, via rapid gene expression regulation [15, 16]. This new knowledge can provide insights into an integrated and dynamic cellular regulatory system whose signal profiles could forecast disease risk associated with early adversity [17–19].

Cells show remarkable flexibility in response to stimuli by regulating gene expression in a transient manner [16, 20]. In one of the first studies relating peripheral blood mononuclear cells (PBMC) gene expression to trauma, basal gene expression signatures, both immediately following trauma and four months later, distinguished survivors who met diagnostic criteria for PTSD from those who did not [21]. Follow-up studies provided further support for associations between chronic stress and glucocorticoid signaling, as well as induced or repressed activation of pro- and anti-inflammatory genes [22–24]. Importantly, several studies have provided evidence of rapid (e.g., from 30 minutes to 8 hours) gene expression activation in response to *in vitro* stimulation [25], psychological stress [26, 27], physical stress [28] and stress-reduction methods [29]. Notably, these response patterns were recently dubbed the “conserved transcriptional response to adversity” [30]. Taken together, theory and evidence suggests that the programmed immune cells of individuals exposed to early adversity may show compromised adaptation in response to acute stress, which, if repeated, may play a downstream role in disease risk.

In this study, we focused on the glucocorticoid-immune signaling pathway by measuring differential expression of glucocorticoid receptor *(NR3C1)*, FK506 binding protein 51 *(FKBP5)* and Nuclear Factor Kappa B Subunit 1 *(NFKB1)* genes [22, 31, 32] in PBMC. The glucocorticoid-immune signaling pathway has been implicated as a key mechanism in relation to chronic stress (i.e., caregiving, poverty [23, 31]), through reduced receptor availability, ligand binding affinity, and functional capacity to regulate gene expression. Specifically, chronic stress, via extended exposure to cortisol, is associated with reduced *NR3C1* expression, leading to glucocorticoid resistance and impaired negative feedback inhibition of the HPA axis [33]. Increased glucocorticoid resistance is additionally explained by increased expression levels of *FKBP5*, an important regulator of the glucocorticoid receptor complex [32]. Reduced levels of glucocorticoid receptors, in turn, bind less cortisol. This effectively decreases the number of ligand-bound receptor complexes available to translocate to the nucleus and regulate the expression of genes, including anti-inflammatory genes. Thus, reduced levels of *NR3C1* expression can lead to impaired immune function. Further, nuclear factor kappa-B (Nf-κB), a highly conserved transcription factor, can increase levels of pro-inflammatory cytokines, partly via reduced inhibition by the ligand-bound glucocorticoid receptor complexes [34]. Evidence exists for elevated levels of pro-inflammatory cytokines in children and adults exposed to ELA [35, 36]. Dysregulation of the immune system in the context of ELA, as well as the resulting increase of pro-inflammatory cytokines, can increase risk for a host of diseases, from autoimmune to atherosclerosis and cancer [37]. Here, in addition to the aforementioned genes, we focused on four pro-inflammatory cytokines, including interleukin-1β (IL-1β), interleukin-6 (IL-6), interleukin-8 (IL-8), and tumor necrosis factor alpha (TNF-α).

We delineate a program of research to study ELA-related programming of biological systems using a within-person, between-groups experimental design. Specifically, in this pilot study we tested whether ELA leads to dysregulation of physiological, gene expression and pro-inflammatory cytokines in response to a canonical laboratory stressor, compared with a resting control condition, and compared with individuals without exposure to ELA. To study the biological embedding of stress, we used a validated screening instrument [38] and recruited 12 men, 6 of whom who endorsed at least 3 significant adverse events (‘ELA group’) [39], and 6 who confirmed zero (‘controls’). In a randomized within-subjects design, we induced acute stress in the lab (Trier Social Stress Test, TSST) and included a no-stress control condition separated by one week. During these sessions, we obtained repeated measurements of physiological reactivity, plasma levels of pro-inflammatory cytokines, and PBMC gene expression over a 4-hour window post-test. We examined how young adults’ exposure to ELA influence neuroendocrine and inflammatory responses to acute stress compared with non-exposed individuals by testing the following: 1) physiological, gene expression, and pro-inflammatory cytokines response to acute stress compared with a no-stress control condition, and 2) stress-induced cortisol changes in gene expression and pro-inflammatory cytokines.

Based on prior literature, we hypothesized that individuals exposed to ELA will evince dysregulated physiological, *NR3C1, FKBP5*, and *NFKB1* gene expression, and pro-inflammatory changes to an acute laboratory stressor, a response pattern that may reveal dynamic biological signatures of early-life programming with implications for life-long health.

## Materials and Methods

### Participants

Participants were healthy male college students at the Pennsylvania State University, recruited by word of mouth and advertisements on campus bulletin boards. We focused on men in this exploratory study due to known sex differences in the stress response [40] and the small sample size for stratified analyses. To obtain the sample who were exposed to ELA, a trained clinical interviewer conducted a phone interview to screen over 100 eligible men using the Stressful Life Events Screening Questionnaire (SLESQ) [38], a 13-item self-report measure that assesses lifetime exposure to traumatic events. We asked respondents 11 specific and two general categories of events, such as death of a parent or sibling, life-threatening accident, and sexual and physical abuse. Based on evidence that three or more traumatic events confers higher risk for disease [39], and considering the severity of the traumatic events, participants who responded to at least 3 incidents up to age 18 years (independently reviewed and reached consensus by MZ and IS) were invited to participate in the ELA group. Respondents’ examples for adverse exposures in this study included (unsubstantiated) child abuse and neglect, severe violence exposure, parental loss, suicide of a close friend or a family member, severe illness of an immediate family member or car accidents. In addition, the SLESQ was used to screen participants without a history of traumatic exposures to serve as the control group. Selection criteria stipulated that subjects were between 18-25 years, without current medical illness or endocrine illness (for example, asthma, diabetes, thyroid disease or pituitary gland disorders confirmed by self-report and physical examination), were currently non-smokers and were not using medication on a regular basis, including psychiatric medication. The final sample included 12 men, 6 of whom experienced early adversity (i.e., ‘ELA group’) and 6 who did not (i.e., ‘controls’) (mean age= 21.25, SD= 2.3). Demographics of the sample are presented in Table 2. The study was approved by the Ethics Committee at the Pennsylvania State University and all participants provided written informed consent. Participants received a modest monetary incentive for participation.

### General Procedure

Testing was carried out at the Pennsylvania State’s Clinical Research Center (CRC). Participants made two visits to the CRC during weekdays, one week apart, on the same day. Testing was scheduled to begin at 11:00am and end by 4:15pm. We used a randomized counter-balanced order for the two sessions (i.e., TSST and no-stress control conditions) blind to participants and lab personnel. Lab personnel were also blind to group status. Participants were given specific instructions to refrain from excessive physical activity on the day of the testing, consuming alcohol for 12 hours before their arrival, and eating and drinking (besides water) for 2 hours prior to the testing session. After arrival and consent, trained nurses completed a physical examination and inserted an IV catheter into the antecubital vein 30 minutes after arrival (30 minutes prior to testing). The TSST session was scheduled to begin at 12:00pm to minimize the effects of circadian changes in cortisol, and was carried out as described previously [41]. Briefly, the TSST consists of a free speech and a mental arithmetic task of 10 minutes duration performed in front of a panel of two committee members (mixed gender) with a camera and microphone situated between the interviewers. Participants were told that they would play the role of an interviewee for a job and have 5 minutes to make an argument for their candidacy. After 5 minutes, the second task emphasizing cognitive load commenced. In this task, participants were asked to count backwards from 1,687 in multiples of 13. If a mistake was made, they were instructed to start again from the beginning. In the no-stress control condition, participants were instructed to sit in a room, read magazines, and to refrain from any stressful activities (e.g., cell-phone use was restricted). After the second blood draw, approximately 60 minutes after the TSST session and 90 minutes after the first baseline measure in the no-stress control condition, participants were administered a set of questionnaires. These questionnaires were administered in both sessions and the average score was calculated before analyses (see below for details). Considering the long time-frame of the study and the repeated collection of multiple blood samples, a standardized low-calorie meal was provided after the third blood draw (approximately at 1:45pm). Fig 1 outlines the study design.

**Fig 1:**
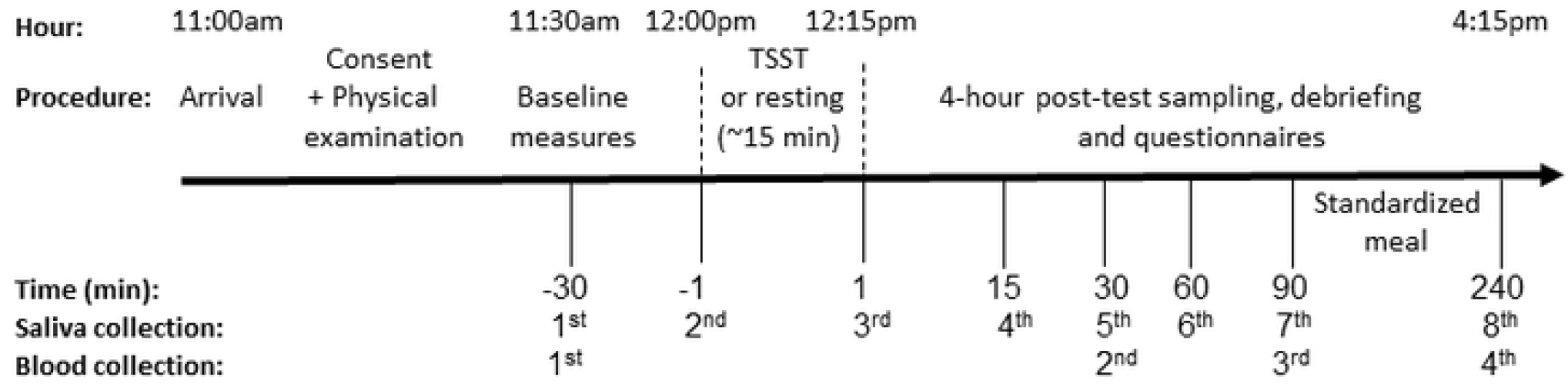
Study design for both sessions (TSST and no-stress control condition), separated by one week

### Physiological Reactivity

Salivary cortisol was repeatedly assessed from the 7 saliva samples at the following time-points: 30 minutes after arrival (30 minutes prior to testing), 1 minutes prior to testing, immediately after testing (15 minutes after last sample in the control condition), and 15, 30, 60 and 90 minutes post-test. Saliva samples were kept at room temperature throughout the session, were immediately centrifuged at the end of the session at 3000 rpm at 24°C for 15 minutes, and then stored at −80°C until assayed. Systolic and diastolic blood-pressure were measured at the same time points as salivary cortisol.

Salivette swabs (Sarstedt, Germany) were used to collect saliva. Salivary cortisol was assessed, in duplicate, through an enzyme immunoassay protocol (Salimetrics) with known controls. The lower detection limit of the assay is <0.007 ug/dL. Intra-assay CV was 9.88% across all samples and inter-assay CV 5.79% across four plates. Participants’ blood pressure was measured, while seated, using an automatic monitor (Omron HEM-712C).

### RNA Extraction and Gene Expression Assays

Gene expression changes were measured repeatedly from the four blood samples at each session at the following time-points: 30 minutes after arrival (30 minutes prior to testing), and at 30 (75 minutes after the first sample in the no-stress condition), 90 and 240 minutes post-test (Fig 1). Given known changes in immune cell redistribution and composition in response to acute stress [20], complete blood count with differential was measured within 24 hours by Quest Diagnostics using additional 4 ml EDTA collection tubes.

Whole blood samples were collected in 10 mL EDTA blood tubes via an IV catheter into the antecubital vein, and immediately centrifuged for 10 minutes at 1500g prior to collection of plasma. PBMCs were immediately isolated through density-gradient centrifugation using Ficoll. Immediately following isolation, cells were suspended in RNAlater solution (Ambion) before being stored at 4 °C overnight. The duration from blood sampling to stabilization of RNA never exceeded 55 minutes. RNA extraction and cDNA synthesis were performed the following day using QIAamp RNA Blood Mini Kit and cDNA Synthesis Kit respectively (Qiagen), and then stored at −80°C until assayed. RNA purity was verified using Nanodrop 2000 spectrophotometer (Thermo Scientific).

All assays were performed on a real-time PCR (Rotor Gene Q, Qiagen). PCR reactions were set-up using the complementary QIAgility robotic pipettor (Qiagen) to ensure maximum pipetting accuracy. Samples were assayed in duplicate. All repeated, within-subject samples were run on the same plate. The reaction mix for gene expression assays consists of 5 uL TaqMan Gene Expression Master Mix (Thermo Fisher Scientific), 1x TaqMan gene expression primer, UltraPure Water (Rockland), and 100ng DNA in a 10 uL reaction. The cycling profile consists of an initial denaturing at 95°C for 15 seconds and annealing/extending at 60°C for 1 minute followed by fluorescence reading, 55 cycles. Three hypothesis-driven genes *(NR3C1:* Hs00353740_m1, *FKBP5:* Hs01561006_m1 and *NFKB1:* Hs00765730_m1) were each normalized to a housekeeping gene *(GADD45A:* Hs00169255_m1). Expression of a given hypothesis-driven gene and the housekeeping gene were assessed on the same plate in two independent PCR reactions using cDNA from the same sample aliquot. Each hypothesis-driven gene was assayed in an independent batch of assays.

Sample normalization was done using the ΔΔCt method [42]. Briefly, a cycle threshold (Ct) is defined as the cycle number at which a sample’s fluorescence reaches a defined threshold. The same threshold was used for reactions assessing housekeeping and hypothesis-driven genes. Thus, each sample on a given plate has two Ct values (e.g. Ct_NR3C1_ and Ct_GADD45A_). The ΔCt is calculated as the difference between the Ct of the gene of interest and the Ct of the housekeeping gene (e.g. ΔCt = Ct_FKBP5_-Ct_GADD45A_). The ΔΔCt represents the within-subject normalization of the three post-test samples to expression levels at baseline. That is, ΔΔCt = ΔCt_POST-TEST_ – ΔCt_BASELINE_. Thus, the ΔΔCt for the baseline sample for each session is always equal to zero. Lastly, fold change is calculated by exponentiating 2 by −ΔΔCt (i.e. Fold Change = 2^-ΔΔCt^). It follows that the fold change for each baseline sample is always equal to one (i.e. 2^0^).

### Pro-Inflammatory Cytokines

Inflammatory assays were performed on plasma isolated from whole blood. Plasma samples were stored at −80°C prior to use. Plasma levels of IL-1β, IL-6, IL-8, and TNF-α were quantified using Meso Scale Discovery’s Multi-Array technology (MSD, V-PLEX Human Proinflammatory Panel II) and analyzed on a Meso QuickPlex SQ 120 instrument (Meso Scale Discovery, Rockville, MD, USA). Sample concentrations were determined relative to standard curves generated by fitting electrochemiluminenscent signal from stock calibrators with known concentrations using MSD Discovery Workbench^®^ software. Samples were run in duplicate. Intra-assay variability was 8.02% across all samples and inter-assay variability was 3.87% across the three plates. The lower limits of detection for inflammatory markers were 0.646 pg/mL (IL-1β), 0.633 pg/mL (IL-6), 0.591 pg/mL (IL-8), and 0.690 pg/mL (TNF-α). Samples with concentrations below the curve fit range were assigned a value of 0 for analyses considering those analytes. This occurred for 13 samples (13.5%) for IL-1β. Samples for all other analytes were within detection ranges.

### Self-Reported Measures and Other Covariates

We administered several questionnaires to assess levels of adverse exposures and mental health symptoms. Specifically, participants completed the following questionnaires at both sessions; the Life-Event Stress Scale (LESS) [43], which consists of 42 common events associated with some degree of disruption of an individual’s life and provide a standardized measure of the impact of a wide range of common stressors; and the Life Events Questionnaire (LEQ) [44], an 82-item inventory-type questionnaire for the measurement of life changes. The LEQ consists of items that are designed primarily for use with students. We further assessed levels of anxiety and depressive symptoms using the Beck anxiety inventory [45], Beck depression inventory [46], and State-Trait Anxiety Inventory [47]), as well as perceived stress levels using the 10-item Perceived Stress Scale [48].

As noted above, given gene expression changes may depend on specific cell populations [20], we measured complete blood cell counts during both experimental sessions, as well as PBMC counts, in duplicate, using a Countess automated cell counter (Invitrogen). Other potential covariates included; age, body mass index, and socioeconomic status (i.e., parental education and income).

### Data Reduction and Final Measures

Statistical analyses of cortisol data used log transformed cortisol values at 7 time-points and area under the curve with respect to increase (AUCi) [49]. The variables were examined for outliers (>3 SD) and none were detected. Blood pressure values were reduced to 4 measures, from 30 minutes prior to testing to 15 minutes after (samples 1-4) to evaluate the fast sympathetic response. Moreover, systolic and diastolic blood pressures were combined to derive a measure of the mean arterial pressure (MAP) to describe the average response in blood pressure (i.e., MAP = [(2 x diastolic) + systolic] / 3). Raw gene expression data was analyzed based on the 2^-(ΔΔCt) method, with normalization to a housekeeping gene, and compared to the first baseline measure in each session [42]. AUCi was computed for each gene to assess overall responses from baseline. Cortisol slope increase was calculated using the first three measures for cortisol from baseline to peak levels and dividing by the time between measures [50].

For the four pro-inflammatory cytokines, considering high correlations [51] (Pearson correlations ranged from .30 to .72), principal component analysis (PCA) indexing *systemic inflammation* of IL-1β, IL-6, IL-8 and TNF-α measures was conducted for the four repeated measures using data from both sessions. In each instance, the first component was extracted for use in subsequent analyses. The four repeated items mapped to components with eigenvalues of 2.55 for the first time point, which explained 63.81% of the variance across all four cytokines, 2.29 for the second time point (57.25% of variance), 2.46 for the third time point (61.54% of variance), and 2.80 for the fourth time point (69.88% of variance). PCA of AUCi for all four cytokines yielded two components with eigenvalues 1.93 and 1.01, which explained 48.27% and 25.37% of the variance respectively. Closer inspection of the factor loading scores for the PCA of AUCi revealed that the first component was largely representative of three cytokines (IL1-β, IL-8, TNF-α) with the second representing IL-6 (Table 1). PCA was also conducted on the four repeated measures independently within each session (TSST and no-stress). The four repeated items in the TSST session mapped onto components with eigenvalues 2.02-2.56, which explained 50.41%-64.01% of variance at each time point. The four repeated items in the no-stress session mapped onto components with eigenvalues 1.65-2.29, which explained 41.28%-57.16% of variance at each time point. The components mapped using data from both sessions were used to investigate within-person differences across sessions, while the components mapped within each session independently were used to investigate between-person differences (i.e. ELA status) within each session. Scores in all repeated questionnaires for both sessions were averaged to increase reliability (Pearson correlations ranged from .72 to .93). None of the demographics measures differed significantly between the ELA and control groups (Table 2), and thus were not included as covariates in the analysis.

**Table 1.**
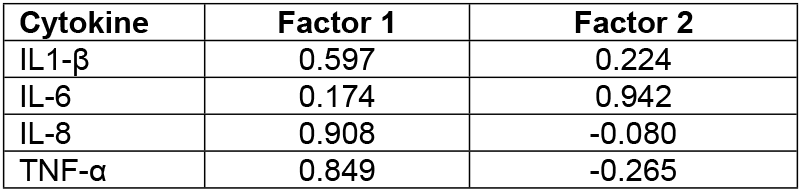
PCA of Pro-Inflammatory Cytokine AUCi: Factor Loading Scores

| Cytokine | Factor 1 | Factor 2 |
| --- | --- | --- |
| IL1-β | 0.597 | 0.224 |
| IL-6 | 0.174 | 0.942 |
| IL-8 | 0.908 | -0.080 |
| TNF-α | 0.849 | -0.265 |

**Table 2.**
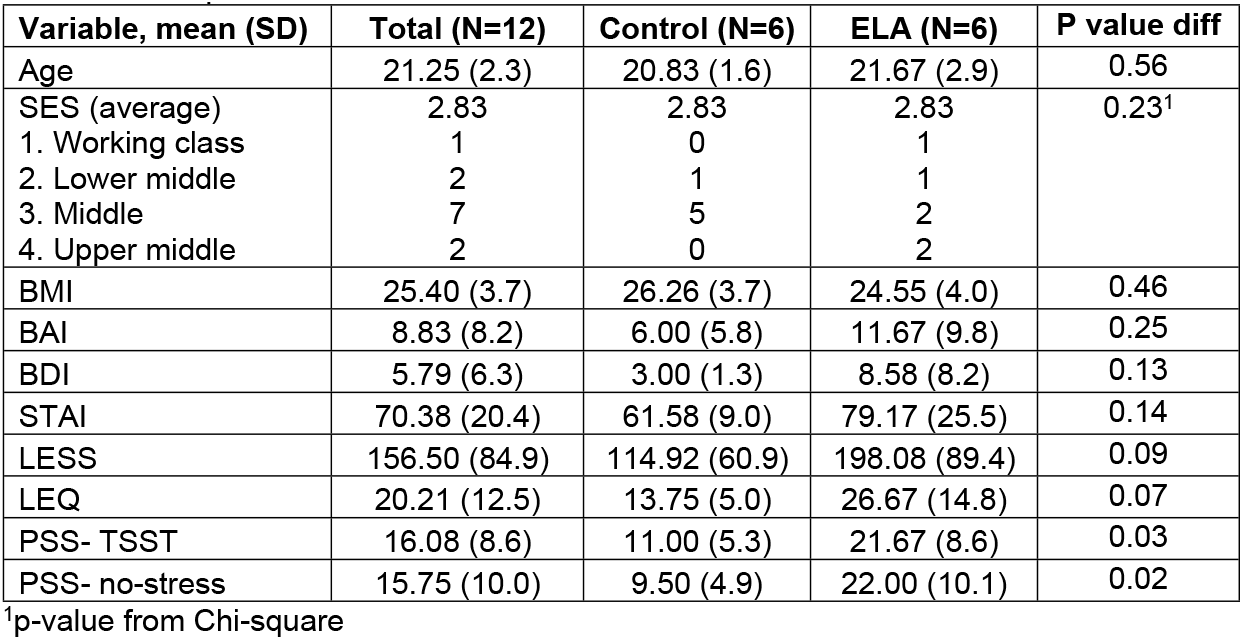
Sample Characteristics

### Statistical Analysis

All statistical tests were carried out using SPSS version 25 (Windows). Repeated measures general linear models (GLMs), ordinary least squares multiple regression analyses, Pearson product-moment correlations, and t-tests were carried out as appropriate. Statistical analyses of changes in gene expression, physiological responses and cytokines levels were subjected to multivariate GLMs, with salivary cortisol, cytokines and gene expression as the repeated measure, condition (stress/no stress) as a within-subjects factor, and status (risk/control) as between-subjects factors. In addition to these analyses, univariate tests were applied to summary cortisol measures (AUCi, [49]) to ascertain reliability of findings, as well as blood pressure (MAP), gene expression, and pro-inflammatory cytokine measures. Huynh-Feldt corrections were applied if sphericity (significant differences in variance between groups) was significant, and only adjusted results are reported.

## Results

### Sample Characteristics and Self-Reported Measures

The ELA and control group did not differ in demographics measures (i.e., age, socioeconomic status and body max index) (Table 2). As expected, the ELA group tended to report more stressful life events [43] compared with controls (univariate ANOVA between-subjects effect: F=3.55, p=0.089 for LESS; F=4.10, p=0.070 for LEQ), as well as higher levels of anxiety [47] (F=2.54, p=0.142), and depressive symptoms [46] (F=2.73, p=0.129). Further, the ELA group self-reported more perceived stress in the TSST session compared with controls (F=6.07, p=0.033), as well as in the no-stress condition (F=7.49, p=0.021), and tended to report more stress in response to the TSST (Likert scale from 1-10) (F=4.02, p=0.080). Overall, these findings confirm previous studies indicating increased stress and anxiety levels in individuals exposed to ELA, compared with non-exposed individuals.

### Physiological, Gene Expression, and Pro-Inflammatory Cytokines Response to Acute Psychosocial Stress Compared with a No-Stress Control Condition

#### Physiological

In the whole sample, repeated measures GLMs indicated significant within-subjects effect for salivary cortisol in response to the TSST, compared with a no-stress condition (Time x Session, F=4.47, p=0.003, estimated effect size η^2^= 0.17), as well as for mean arterial pressure (Time x Session, F=5.31, p=0.003, η^2^= 0.20). Compared with controls, the ELA group exhibited significantly higher mean arterial pressure response to the TSST (Time x Status, F=8.59, p<0.001, η^2^= 0.46), and a trend towards a higher cortisol response in the TSST relative to no stress (ΔAUCi: F=3.58, p=0.088) (Fig 2). Notably, no significant differences were observed between the ELA and control groups in the no-stress condition (Time x Status, F=1.38, p=0.257 for salivary cortisol; F=1.01, p=0.402 for MAP). Overall, these findings confirm some [52], but not all studies [7], indicating increased physiological reactivity to acute stress in young adults exposed to early adversity, compared with non-exposed individuals.

**Fig 2:**
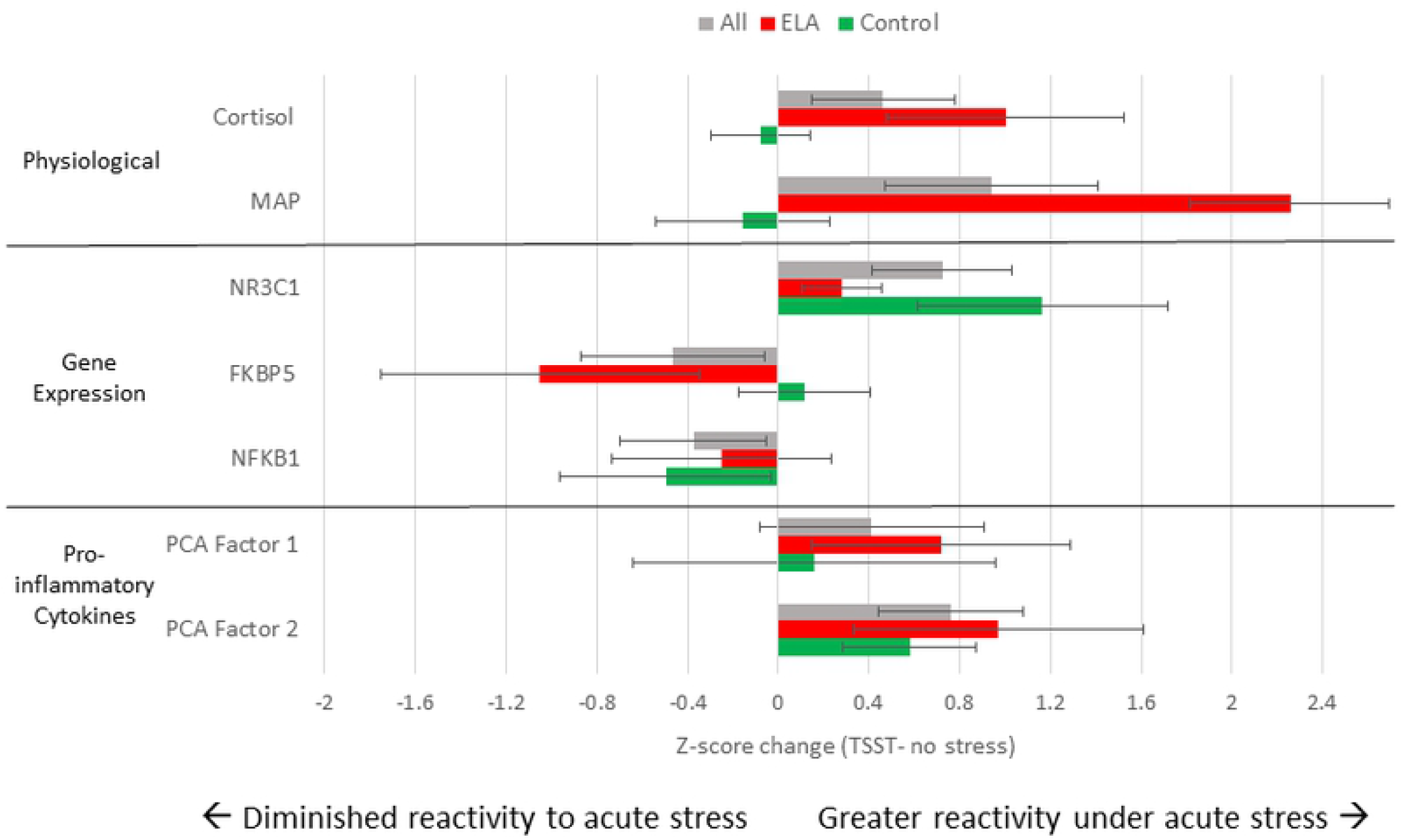
Normalized change score for physiological, gene expression, and pro-inflammatory cytokine response to the TSST relative to the no-stress session for ELA group, control group and full sample. Change scores were calculated by standardizing summary AUCi using data from both sessions and subtracting participant values from the no-stress session from those in the TSST session. Error bars represent standard error of the mean. Change scores are expressed for the full sample (grey), ELA group (red), and control group (green).

#### Gene Expression

In the whole sample, there was a significant within-subjects effect of TSST vs. no-stress condition on *NR3C1* gene expression with increased levels in the TSST (Time x Session, F=4.85, p=0.006, estimated effect size η^2^= 0.19). Group analysis revealed increased levels in the control group (Time x Session, F=5.09, p=0.013, estimated effect size η^2^= 0.36), but not in the ELA group, which had a blunted response to the TSST (Time x Session, F=1.00, p=0.406, estimated effect size η^2^= 0.09) (Fig 3A), suggestive of *NR3C1* expression resistance and lower levels to inhibit the HPA axis. Notably, no differences were observed in *NR3C1* expression between the ELA and control groups in the no-stress condition (Time x Session, F=0.78, p=0.491) (Fig 3B). *FKBP5* and *NFKB1* expression did not change significantly in response to the TSST vs. no-stress condition, and responses did not differ by group status.

**Fig 3:**
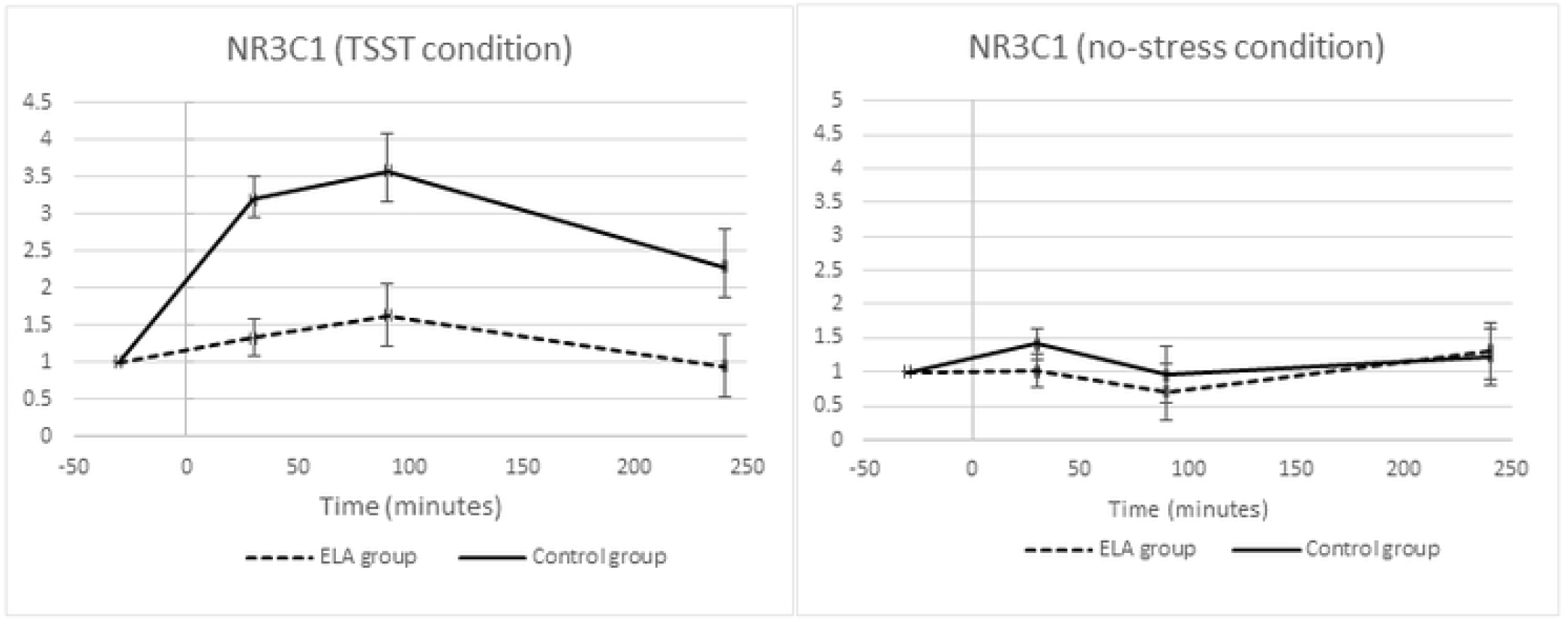
Fold change in NR3C1 for ELA (dashed lines) and control groups (solid lines) in response to the TSST (left) and during the no-stress sessions (right). Error bars represent standard error of the mean.

#### Pro-Inflammatory Cytokines

In the whole sample, PCA for the four repeated measures of IL-1β, IL-6, IL-8 and TNF-α did not reveal a significant within-subjects effect of TSST vs. no-stress condition using repeated measures GLM analysis (Time x Session, F=0.29, p=0.831). Similarly, an analysis of the first AUCi component did not reveal significant differences in systemic inflammation between the TSST and no-stress conditions (F=2.09, p=0.165). However, an analysis of the second AUCi component (largely representing IL-6) showed significantly greater pro-inflammatory responses to the TSST relative to the no-stress condition (F=5.85, p=0.026) (Fig 2). No differences were observed between the ELA and control groups in response to the TSST relative to no-stress using either AUCi PCA components.

Exploratory analyses of each pro-inflammatory cytokine revealed a significant within-subjects effect for IL-6 in response to the TSST (Time x Session, F=2.97, p=0.044, AUCi: F=7.70, p=0.018), but not for the other three cytokines (IL-1β, Time x Session, F=0.80, p=0.500, AUCi, F=1.37, p=0.257; IL-8, Time x Session, F=0.85, p=0.470, AUCi, F=0.93, p=0.434; TNF-α, Time x Session, F=0.25, p=0.863, AUCi, F=0.40, p=0.537). Again, responses did not differ by group status.

### Stress-Induced Cortisol Changes in Gene Expression and Pro-Inflammatory Cytokines

Ordinary least squares multiple regression analyses tested whether stress-induced cortisol increase in response to the TSST predicted changes in gene expression and cytokines, and whether the responses differ between the ELA and control groups. Specifically, we used cortisol slope increase from baseline to peak levels (from 30 minutes prior to testing to 15 minutes after stress onset) to predict summary changes in gene expression and cytokines in response to the TSST (Fig 4).

**Fig 4:**
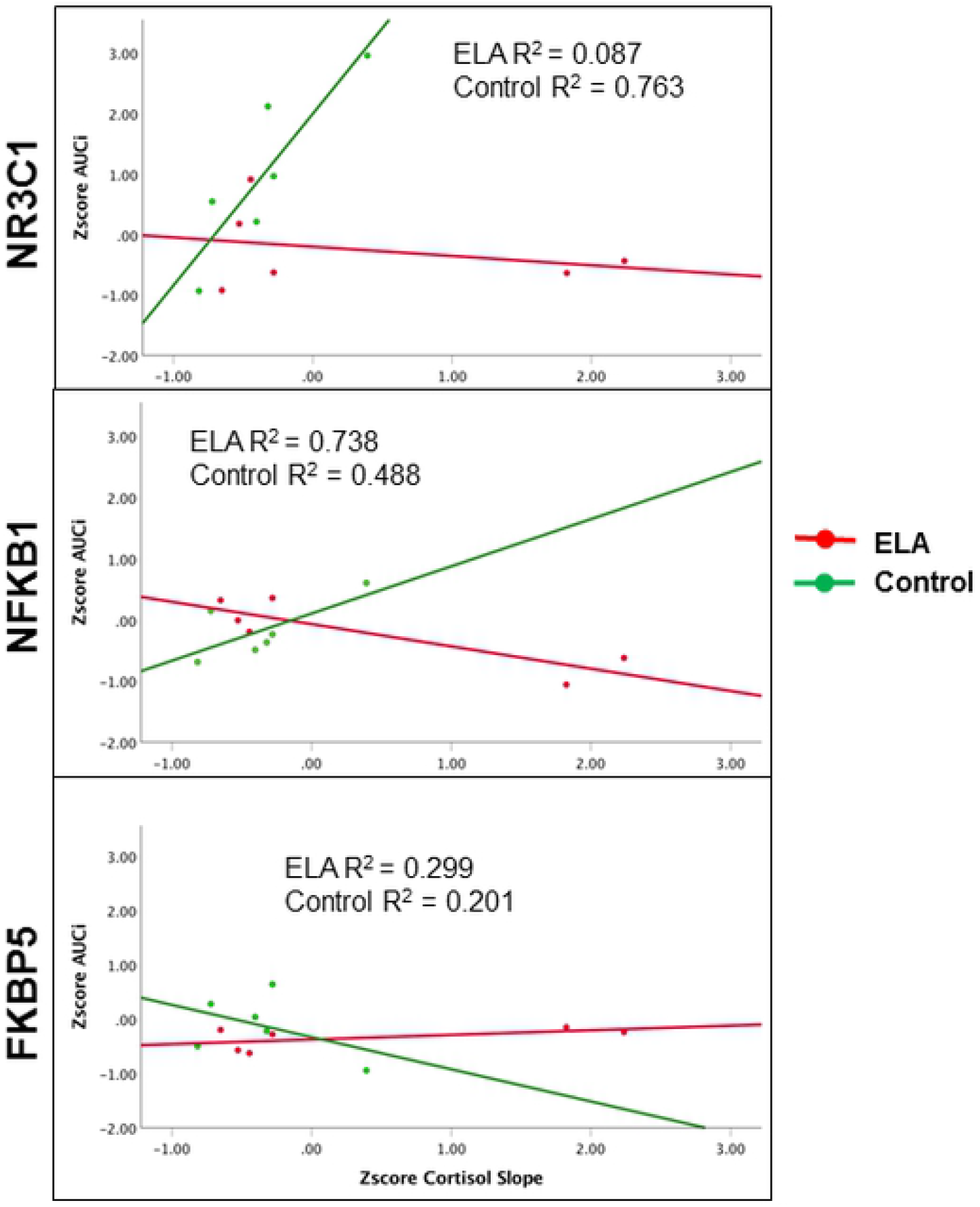
Scatterplot and fit lines for summary gene expression changes in NR3C1 (top), NFKB1 (middle), and FKBP5 (bottom) in response to stress induced cortisol increase for ELA group (red) and control group (green). Cortisol increase calculated as the slope from baseline to peak levels, and then standardized for figure construction. Gene expression changes expressed as AUCi summary measure, which was then standardized for ease of comparison across genes. R^2^ shown are from models with cortisol slope as the only predictor.

In the whole sample, cortisol increase did not predict significant changes in gene expression over time (p= 0.728 for *NR3C1;* p= 0.156 for *NFKB1*, p= 0.832 for *FKBP5)*. When group status was included in the regression analyses, cortisol increase predicted significant changes in *NR3C1* and *NFKB1* gene expression in response to the TSST (β= −2.66, t= −2.90, p=0.023 for *NR3C1;* β= −3.61, t= −3.67, p=0.008 for *NFKB1)*. Moreover, cortisol increase interacted with group status such that increase in cortisol predicted increase in both *NR3C1* and *NFKB1* expression among controls, but decrease in the ELA group *(NR3C1*, Group x Cortisol Increase β= 2.74, t= 3.35, p=0.012; *NFKB1*, Group x Cortisol Increase β= 2.86, t= 3.25, p=0.014). For *FKBP5*, group status and cortisol interaction did not reach statistical significance (Group x Cortisol Increase β= −1.84, t= −1.36, p=0.215).

For pro-inflammatory cytokines, in the whole sample, cortisol increase did not predict significant changes in cytokines using a PCA for the repeated measures (β= −0.12, t= −0.36, p=0.730). Further, there was no interaction by group status (Group x Cortisol Increase, β= 1.00, t= 0.67, p=0.525) (Fig 4).

### Sensitivity Analyses

Sensitivity analyses were conducted using the ‘leave-one-out’ method. Overall, results were robust to the removal of any individual participant. Differences in sample characteristics and self-report measures remained consistent upon removal of any given participant, as did physiological, inflammatory, and gene expression responses to the TSST relative to the no-stress session. Likewise, the ELA group continued to display increased MAP responses to the TSST relative to the control group. Differences between ELA and control groups in cortisol response to the TSST relative to no-stress (ΔAUCi) were modestly attenuated by removal of any given participant, but not appear to be driven by a single individual.

Removal of one participant in the ELA group did modify associations between stress-induced cortisol changes and *NR3C1* and *NFKB1* gene expression. Specifically, cortisol increase and group status no longer interacted to predict gene expression over time (Group x Cortisol Increase, β= 1.637, t= 0.626, p=0.554 for *NR3C1;* β= 1.537, t= 0.474, p=0.652 for *NFKB1)*. Instead, removal of this participant increased the contribution of group status in the model, such that both cortisol slope and group status were independently associated with gene expression changes, without an interactive effect (Group Status, β= 3.524, t=3.671, p=0.008 for *NR3C1;* β= 2.845, t=2.418, p=0.046 for *NFKB1)*. By contrast, removal of a different participant from the control group resulted in associations between stress induced cortisol increase and *FKBP5* gene expression that were previously unobserved. Specifically, models run without this participant showed a significant association between cortisol increase and *FKBP5* gene expression (β= 3.177, t=2.534, p=0.044) as well as an interactive effect between group status and cortisol slope (Group x Cortisol Increase, β= −3.114, t= −2.750, p=0.033).

## Discussion

To our knowledge, this is the first investigation of stress-induced gene expression and pro-inflammatory cytokines changes within-individuals, comparing stratified groups of ELA-exposed and control individuals. By comparing a validated laboratory-based stressor to a no-stress condition within the same individuals, we were able to disentangle the effects of acute stress from noisy measurements in the same individuals. Further, this design allowed us to distinctly identify if/when differences between ELA-exposed and control individuals were context dependent (i.e. manifesting only during stress). Results provide preliminary evidence in humans of a dysregulated pattern of *NR3C1, FKBP5* and *NFKB1* gene expression activation as a consequence of ELA. Importantly, these changes manifest more acutely in the presence of stress-induced cortisol release as compared to a no-stress resting condition.

As predicted by previous research, the ELA group evince higher cortisol response and lower *NR3C1* gene expression in response to the TSST compared with controls, with no difference between groups in the no-stress condition. Moreover, cortisol-induced changes in gene expression revealed a decoupling between the stress-induced cortisol release and nuclear signaling in the ELA group. Cortisol reactivity was associated with increased *NR3C1* and *NFKB1* expression in the control group, but in the ELA group these associations were blunted. Findings for cortisol-induced *FKBP5* expression revealed the hypothesized pattern of increased activation in the ELA group, and decrease among control individuals, although results did not reach statistical significance in the full sample. For pro-inflammatory cytokines, only IL-6 increased significantly in response to the laboratory-induced stressor, however, stress-induced cortisol release did not predict changes in cytokines levels, contrary to hypothesized prediction. Overall, we provide preliminary findings for the biological embedding of ELA via a dynamic and dysregulated pattern spanning multiple levels of analysis (genomic and physiological), and which presents more acutely in response to psychosocial stress.

Findings concur with the receptor-mediated model of glucocorticoid signaling resistance [53]. First, ELA was associated with increased cortisol response to the TSST compared with controls, confirming some [52], but not all studies [7], indicating increased physiological reactivity in young adults exposed to early adversity. Second, chronic exposure to cortisol, as a consequence of ELA, can lead to a compensatory response whereby glucocorticoid sensitivity decreases (e.g. via decreased receptor availability). Here we replicated prior evidence of reduced *NR3C1* expression levels in ELA-exposed individuals, but only in response to acute laboratory stress. We also provide preliminary evidence that the reduced *NR3C1* expression is driven by decreased responsiveness to stress-induced cortisol release into the periphery. Third, FKBP5 has been implicated in the glucocorticoid resistance model whereby overexpression of FKBP5 reduces cortisol binding affinity to glucocorticoid receptors and further translocation to the nucleus [54]. Here, we do not confirm previous findings. Future studies employing larger sample sizes may be required to test for *FKBP5* response as these effects may be more subtle and/or sensitive to outliers (i.e. sensitivity analyses without a given control participant were in line with predictions). Fourth, diminished availability of the ligand-bound glucocorticoid receptor complexes in immune cells is suggested to contribute to reduced inhibition of Nf-κB signaling, leading to increased pro-inflammatory cytokines [34]. Here, we tested whether ELA is associated with increased activation of *NFKB1* gene, a DNA binding subunit of the Nf-κB protein complex. In line with expectations, stress-induced cortisol increase was associated with increased *NFKB1* expression among controls, suggesting decreased inhibitory action on Nf-κB signaling [55]. In the ELA group, however, stress-induced cortisol increase was associated with decrease *NFKB1* expression. Fifth, in vitro studies have established a connection between glucocorticoid exposure and diminished capacity of immune cells to inhibit pro-inflammatory cytokines in individuals exposed to psychological stress. Here, only pro-inflammatory cytokine IL-6 increased significantly in response to acute stress, replicating previous studies [56]. However, contrary to expectation, stress-induced cortisol release did not predict increased pro-inflammatory profile among individuals exposed to ELA.

The methodological strengths of this study include a laboratory-based within-subjects experimental design, which allows stronger causal inferences. We collected repeated measurements over a relatively long time scale to document changes in gene expression and pro-inflammatory cytokines. Our within-subjects, between-groups design, combined with four repeated measurements in each session, reduced biological variability and increased power to detect true associations. Finally, we tested the moderating effects of ELA, which enables tests of potential programming of biological systems.

We acknowledge limitations. First, this was a pilot study with a small sample size. Although comparable to similar prior investigations [26, 31, 57], the results from this study still need to be interpreted with caution. Notwithstanding, the strength of the within-subjects experimental design combined with the leave-one-out sensitivity analysis alleviate concerns about spurious findings. We focused on men in this exploratory study due to known sex differences in the stress response [40] and the small sample size for stratified analyses. Future studies with larger sample size including both males and females are needed to replicate these findings. Second, gene expression changes are tissue-specific. As a first test, we isolated PBMC from whole blood to measure gene expression changes. The exclusion of granulocytes cells provides a cleaner measure of the more active populations of lymphocytes and monocytes. Nevertheless, future studies will benefit by measuring gene expression changes in specific sub-populations of leukocytes. Third, we focused on three hypothesis-driven genes. There are multiple biological pathways that are activated in response to stress that may play a downstream role in disease susceptibility, such as the conserved transcriptional response to adversity pathway [30]. Prior research has investigated multiple genes using microarrays [23, 26–28, 31]. Future studies with adequate sample size will benefit by testing larger groups of genes/pathways. Fourth, this study did not consider specific types of ELA, or timing of exposure. Here, we focused specifically on severity of multiple (i.e., minimum of three) ELA exposures up to age 18 years. Future research can explore specific types of ELA in different populations and settings. Further, our study included non-Hispanic white males and thus future research need to test whether the association generalizes to other populations. Finally, although we included a no-stress condition to control for the higher degree of noise associated with gene expression measurements, the control session did include the stress of venipuncture. However, this is unavoidable technical limitation for collecting sufficient immune cells for gene expression research.

In conclusion, ELA may program physiological systems in a maladaptive manner more likely to manifest during times of duress, predisposing individuals to the negative health consequences of everyday stressors. Although increased activation of the glucocorticoid-immune signaling in response to acute stress is considered adaptive in the short-term, persistent activation can increase risk for mental and physical health problems. These results could potentially identify new targets for therapeutic interventions mitigating the negative effects of early adversity, such as pharmacological agents acting on the glucocorticoid receptor and FKBP5 [32]. Further, while previous risk factors and biomarkers of stress contributed to our understanding of biological embedding processes, these are nevertheless static characteristics that have not explained health outcomes very well. For example, considering high failure rates for depression treatments, and in order to tailor individual interventions, identifying objective changes in stress-induced gene expression may help to predict short-term intervention efficacy in clinical and non-clinical settings. An example for such an effort could be to utilize models of dynamic cellular markers as individual-level factors to account for variation in intervention response and clinical outcomes [17–19]. Thus, future research in this area can have a range of impacts for basic science, intervention studies and clinical practice that will influence treatments to match the specific cellular processes operating within an individual.

## Acknowledgments

We thank all nurses at the CRC and the participants in this study. Research reported in this publication was supported by the National Institutes of Health, National Institutes of Aging through R21AG055621 grant (I.S.) and by the National Center for Advancing Translational Sciences through UL1 TR002014 grant. W.J.H. and L.E. were supported by National Institute on Aging T32 AG049676 to The Pennsylvania State University. The content is solely the responsibility of the authors and does not necessarily represent the official views of the National Institutes of Health.

